# Statistics used by ecologists: the rise of R statistical software, GLM(M)s and Networks

**DOI:** 10.64898/2025.12.04.692382

**Authors:** Estevao Alves da Silva

## Abstract

About 15 years ago, statistical softwares were rarely employed to conduct statistical analyses, and biologists/ecologists used roughly 100 statistical procedures in research. This number has likely increased substantially with the development of new analytical methods and the widespread adoption of computers and softwares. In this study, I investigated the temporal variation in statistical procedures and the use of softwares used in studies on plant–ant interactions. Data were collected from 142 published papers covering a period between 1979 to 2023. Information related to statistics terminology, the softwares cited and R software packages were retrieved. Each paper had on average eight statistical procedures and there was a significant increase in procedures over the years. The R software was cited in almost 80% of studies, surpassing by far the other softwares. Classical analyses such as *t*-tests, correlations and chi-squares are still used in research; nonetheless, from 2012 onward a new set of analyses including GLM, GLMM and Networks have emerged and are being used as the leading analyses in studies of plant-ant interactions. This coincides with the adoption of R software by scientists in this field. The most used R packages were *vegan, lme4* and *bipartite*, followed by several other related to linear (mixed) models. Statistics literacy is a mandatory aspect of research in plant-ant interactions. The increase in statistical procedures over the years, as well as the emergence of new tests can be considered as a natural evolution of the field with demand for robustness in methods and analyses.

## Introduction

About 15 years ago Velásquez et al., (2011) demonstrated that that *t*-tests, Anova, correlations, chi-squared tests, and regressions were among the most frequently used statistical procedures used by biologists. Due to their simplicity, these classical tests could be performed without programming or specialized software (Hobbs and Hilborn, 2006; McCrum-Gardner, 2008; Moser and Stevens, 1992). However, the widespread availability of computers and advanced statistical softwares over the past decade has expanded the range of analytical methods in biological and ecological research. It is not uncommon to encounter papers in which ecologists use a wide variety of tests, with some studies employing tens of different analyses of data (Miranda et al., 2019). Notably, there has been a substantial increase in the application of generalized linear models (GLMs), generalized linear mixed models (GLMMs), and Network analyses, particularly in studies addressing plant–ant interactions (Dáttilo et al., 2013; Sendoya et al., 2016; Souza et al., 2024). These modern analytical approaches are designed to uncover new trends and patterns in ecological data which would not be possible, or at least inappropriate by using classical tests (Bolker, 2015).

Modern statistical methods also require specialized software, with the R statistical environment (R Core Team, 2025) being the apparent preferred choice among ecologists (Gao et al., 2025). Indeed, many tests are developed to run almost exclusively in R or perform better within this program (Jaeger et al., 2017; Knudson et al., 2021; Luke, 2015). R essentially requires a basic understanding of programming, has a steep learning curve, and is known for its non-intuitive interface, which differs markedly from point-and-click softwares that offer accessible toolbars (Culpepper and Aguinis, 2011; Davidson et al., 2019). In R, scientists must not only be familiar with the terminology inherent to statistics, but also be able to translate this knowledge into programming commands (Woodard and Lee, 2021).

R is a comprehensive software environment, allowing users to manage datasets (whether stored locally or accessed online), conduct statistical analyses and generate figures. This versatility is possible because in addition to its base functions, R provides access to a vast repository of thousands of packages (Lai et al., 2019) that can be added to the software to largely expand its analytical capabilities (Lortie et al., 2020). Indeed, several analyses commonly used by ecologists, such as GLMMs and Network analyses, cannot be performed without specialized packages (Brooks et al., 2017; Dáttilo and Rico-Gray, 2018). In this context, the R packages cited in papers not only reflect the type of analyses conducted by scientists, but also the type and complexity of data under investigation.

The array of analyses possible to perform in R and its packages, combined with study designs that accommodate the collection of large datasets, has presumably increased the amount of data analyses presented in scientific papers. Thus, in this study, I investigated the temporal variation in statistical procedures and data analyses used in ecological research, specifically in studies on plant–ant interactions. To simplify the text, the expression “*statistical procedure*” will be used hereafter to refer to the statistical tests, quantitative analyses, indices and nomenclatures related to data analysis.

The following questions were addressed: *(i)* Has the number of statistical procedures increased over the years, and was it related to the adoption of softwares? *(ii)* How often do ecologists use R software and its packages? *(iii)* How similar are the packages used in studies of plant-ant interactions with other fields of ecological studies? (e.g., Lai et al., 2019; 2023a; 2023b) *(iv)* Is there a temporal variation in the use of traditional (e.g., *t*-tests, correlations) and modern tests (GLMs, GLMMs and Networks) in ecological research?

## Methods

### Data collection - plant-ant interactions

The statistical procedures used in this study were retrieved from the literature on plant–ant interactions, particularly research on plants bearing extrafloral nectaries. Studies on this subject, whether observational or experimental, have explored a wide range of topics (Cardoso et al., 2023; Moura and Del-Claro, 2023; Nogueira et al., 2015), thus allowing a compilation of a diverse set of statistical procedures applied in various contexts. This approach also provides a suitable sample size for comparison and enables examination of temporal variations in the use of statistics and softwares.

It is important to note that the statistical procedures presented hereafter are not specific to plant–ant studies. In fact, most are widely used across different disciplines (Bettany-Saltikov and Whittaker, 2014; Mangiafico, 2015; Paquot and Th.Gries, 2020), while others have been adapted from non-ecological fields and implemented in ecological research (Hevey, 2018; Newman, 2002).

The literature used in this research was obtained from Barbosa et al., (2025) who compiled a list of studies on plant–ant interactions (up until 2022) mediated by extrafloral nectar in Brazil. To supplement the dataset, an additional literature search was conducted between September and November of 2024 using Google Scholar. The search terms included *extrafloral*, *ants*, and *Brazil* and I restricted the search to papers published until 2023. A series of papers was downloaded and cross-referenced with Barbosa et al. (2025) to remove duplicates. Subsequently, the remaining papers underwent detailed examination and thorough reading. Unlike Barbosa et al. (2025), I considered plant–ant interactions in *ex situ* conditions (e.g., in gardens, greenhouses, and botanical gardens). This approach not only could increase the sample size but also includes statistical procedures used in different study designs and situations.

Only papers published in journals with editorial boards, written in English (as a measure to standardize the nomenclature of statistical terms), and with at least one analysis of data were included in research. Data extracted from each paper included the authorship, year of publication, statistical procedures used, softwares employed, and R packages cited.

### Identification of statistical procedures

The statistical nomenclature was extracted from the Methods and Results sections of each paper in their original form, as described by authors. Every terminology which was related to statistics was retrieved (except for descriptive statistical terms such as mean, sd, median, etc.); so, the data contains a wide array of terms, from simple analyses like transformations, overdispersions and normality checking; to more specific tests such as Circular Statistics, GLM(M), and the full collection of Networks indices (Online Resource). In short, all terms indicative of statistical procedures were registered.

I then compiled a list with standardized names for each procedure. To accomplish this, I examined each statistical procedure described in papers and provided a name indicative of the test. I found hundreds of terms/names related to statistical procedures, but many of them were synonymous. For example, linear regression was described as *regression, linear model, simple linear regression*, or *simple regression*. This issue was also noted by Velásquez et al. (2011). Several adaptations were made to standardize the test‘s names and I used the best of my knowledge and experience in statistics to relate each statistical term to its actual test. In case of doubt, I consulted the literature and the source material.

From each single paper in which data was retrieved, I considered only unique names of statistical procedures, and excluded repetitions (e.g., the same procedure that was described consistently in the Methods, such as transformation, overdispersion, null model), plurals (e.g., *t*-test and *t*-tests) and synonyms, which eventually refer to the same test (e.g., *t*-test and Student’s *t*-test). In addition, each distribution error of GLMs and GLMMs (e.g., Poisson, negative binomial, etc.) was considered as a different statistical procedure; if the distribution error was not provided, these tests were regarded as GLM or GLMM only. All types of Anova (one-way, two-way, repeated measures, etc.) and *t*-tests (one sample, two sample, paired) were also considered different procedures. Conversely, other procedures were grouped into one single name, such as chi-square and chi-square with Yattes correction; the different forms of Akaike Information Criterion (weighted, for small samples and corrected) and the NODF index (weighted or unweighted nestedness based on overlap and decreasing fill). After all papers were read and analyzed, I found a total of 784 terms related to statistical procedures, which were then reduced to 261 standardized names (Online Resource).

### Traditional and modern analyses

To investigate the temporal variation of traditional and modern statistical analyses used in plant-ant interaction studies, I selected the most frequent tests (> 10% of use in literature) (Online resource) which were leading analyses and related to hypotheses testing. The traditional tests include seven tests, namely the Student‘s *t* test, Mann-Whitney *U* test, Chi-square test, linear regression, one-way Anova, Kruskal-Wallis test and the paired Student‘s *t* test. The modern tests comprise the GLMs (all error distributions), GLMMs (all error distributions) and Network analyses (Online resource 2).

These tests can be classified as classic/traditional and modern according to literature (Bettany-Saltikov and Whittaker, 2014; Bolker, 2015; Brezina, 2020; Heit et al., 2024; Hobbs and Hilborn, 2006; Lehmann, 2011; Rutherford, 2011; Thompson, 2013). The so-called modern tests have been implemented in ecology recently (Bolker, 2015; Dáttilo and Rico-Gray, 2018; Pekár and Brabec, 2016); in contrast, the traditional tests are usually taught in statistics classes and textbooks, and some are over 100 years old (Bzdok, 2017). It is important to note that Networks are not single tests *per se* but rather analytical procedures which involve multiple analyses and provide numerous metrics and indices (Dáttilo and Rico-Gray, 2018). This approach also allows the exploration of modern methods in relation to the adoption of statistical softwares, particularly R. Papers were classified as having each of these groups of tests.

### Softwares and R packages

All softwares used by researchers were listed and no distinction was made between those used for statistical analyses and those used for figure generation, as researchers might not specify the purpose of each software. Softwares’ names are presented without version numbers, and RStudio was classified as R software.

From each paper that cited R software, I investigated the Methods and References sections to retrieve the information of the packages used for analyses.

### Statistical analysis

A generalized linear model (GLM) with a negative binomial distribution was used to investigate whether the number of statistical procedures per paper was related to years (continuous variable) and to the softwares used (categorical factor) (***objective i***). In this analysis I used the data as described in subsection *Identification of statistical procedures*, which yielded a sample size of 261 statistical procedures. Softwares were classified into four categories: R software (papers that used R exclusively), R+others (papers that used R in combination with other software), others (which included several softwares, but with low frequency to justify their individual inclusion), and unknown (papers where softwares were not cited). The choice of using R software as a categorical variable in this analysis is justified because R was present in most publications (see Results).

Data of use of R software and its packages (***objective ii***) are shown in figures, which depicts the frequency of papers that cite R (in relation to all softwares used) and the temporal variation in the use of R by ecologists. One figure also shows the most used R packages in research, and to accommodate a number of packages, I show only those which were present in three papers or more. No statistical test was made in this approach. The increase in the use of R software over the years (as also examined in Gao et al. 2025) was examined with a linear regression. The dependent variable was the frequency (in % values) of papers per year that cited R and the independent variable was years, specifically from 2012 onwards, which is period in which R was implemented in plant-ant studies.

My data on the most frequent packages used in plant-ant studies were compared with the most used R packages in ecology, biodiversity conservation and forestry (Figure 2 in Lai et al., 2019; Figure 3 in 2023a; Figure 3 in 2023b) (***objective iii***). I used *n* = 16 packages for comparison among works, as this is the threshold I used to show the most frequent packages in plant-ant studies (see Results section). For this comparison I used the Jaccard index of dissimilarity. I only compared the top-used packages between my work and Lai‘s et al. studies because the simple sizes of the works were very different, being inappropriate to compare all packages used.

**Fig 1.**
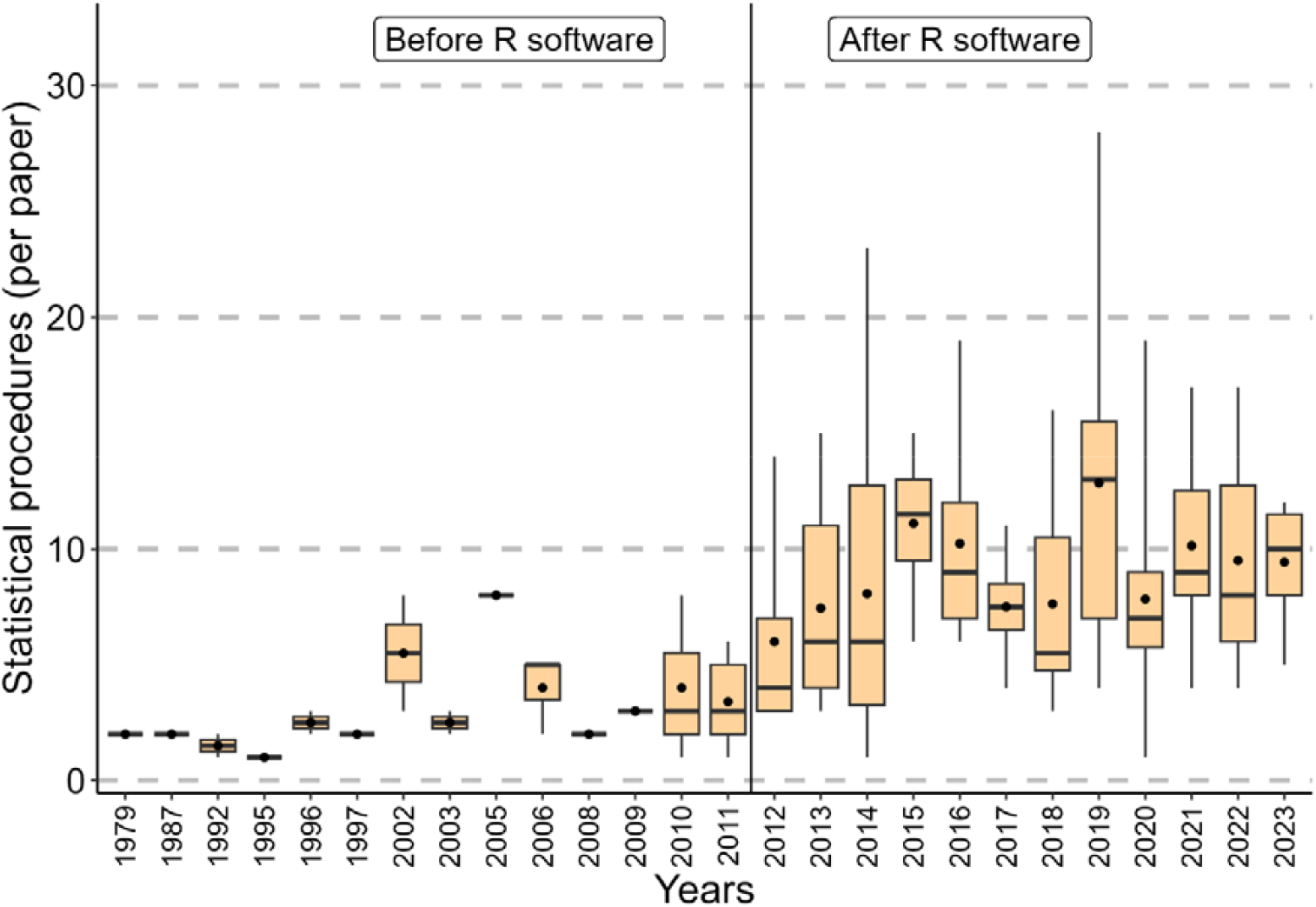
Temporal variation in the use of statistical procedures in plant-ant interactions. The number of statistical procedures increased from 2012 onward, and it was related to the adoption of R software. The circle inside boxes indicates the mean value

**Fig 2.**
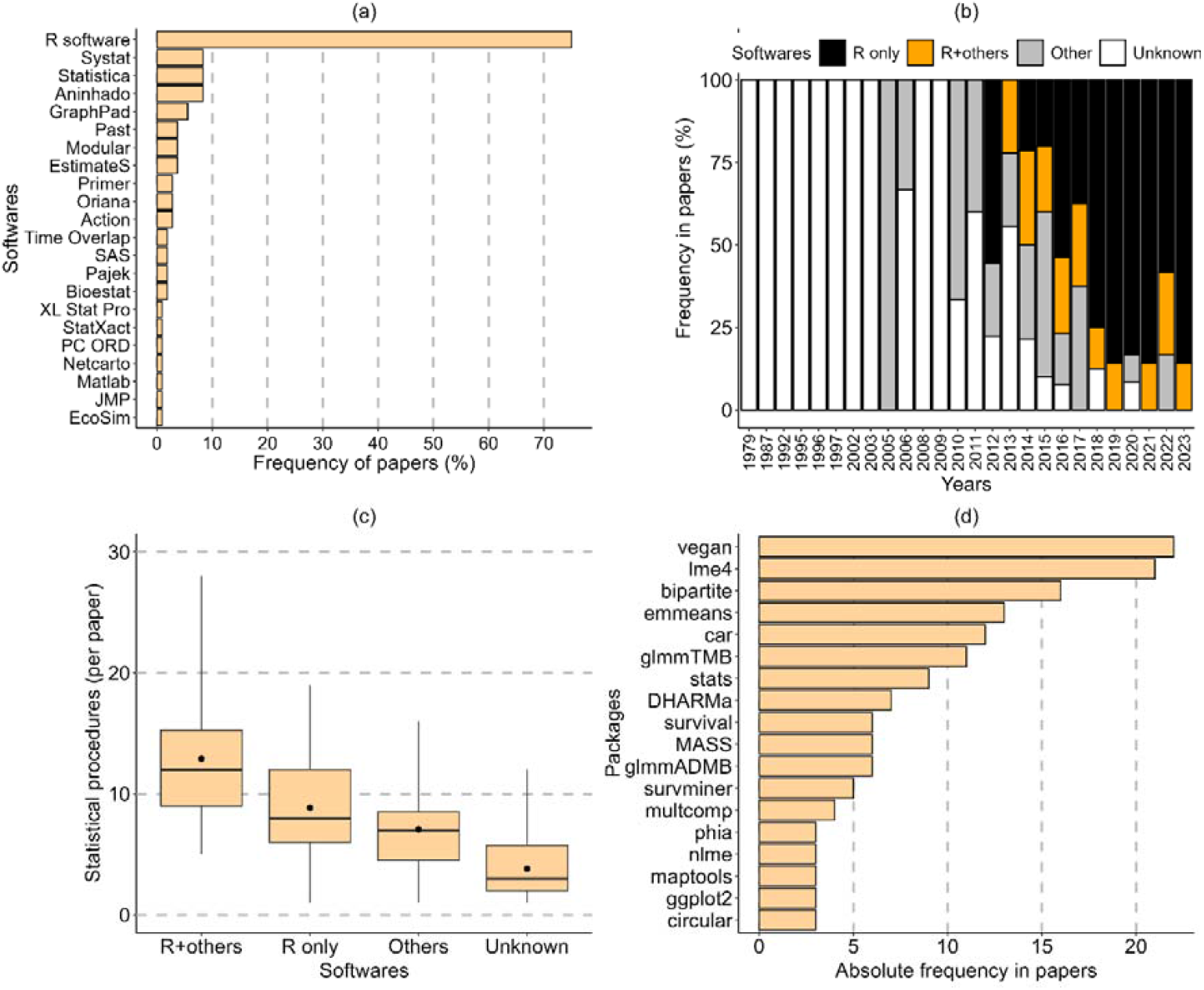
**(a)** Frequency of softwares used in papers. **(b)** Use of R software by scientists over time. **(c)** Number of statistical procedures per paper according to softwares. “R+others” (*n* = 20 papers) indicates that R was used together with other softwares; “R only” (*n* = 61 papers) means that only R was used; “Others” (*n* = 27 papers) refer to softwares showed in **“a”** that were also used in papers (except for R). “Unknown” means that papers did not cite the use of softwares (*n* = 34 papers). The circle inside boxes in **“c”** indicates the mean value. (**d**) Most frequent R packages used in plant-ant studies.

**Fig 3.**
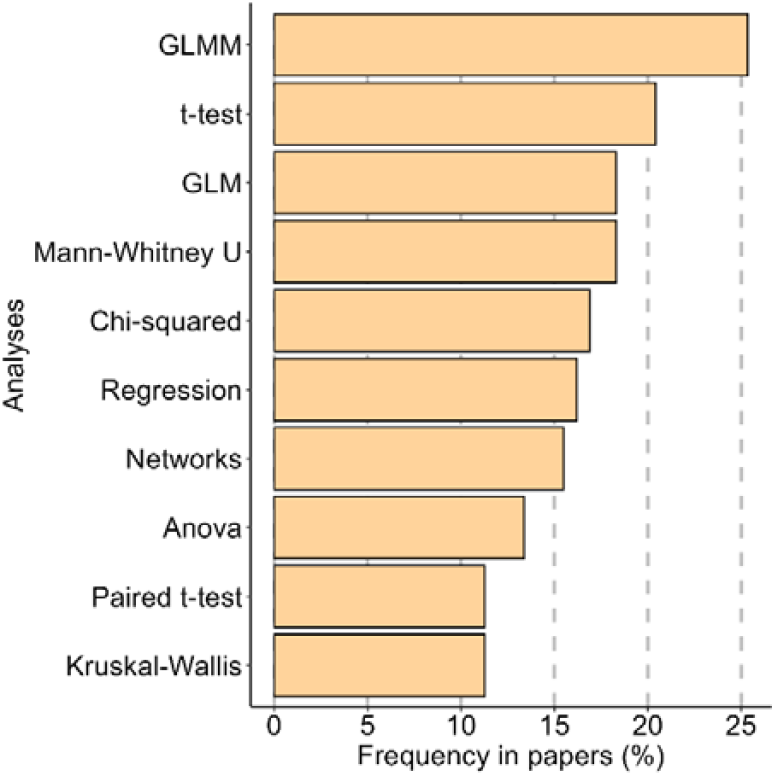
Frequency of different analyses in papers dealing with plant-ant interactions.

The variation in the use of traditional and modern analyses (subsection *Traditional and modern analyses*) (***objective iv***) is shown in a figure that displays how many times these tests appeared in papers over the years. No statistical test was used in this comparison. Data were not independent since many papers employed multiple analyses. For example, a paper using a *t*-test might also include a GLMM or other tests. In addition, data contained many zero values, as some tests have been implemented in research only in recent years. All statistical analyses and figures were made using R software.

### Results

The literature search accounted for 142 papers, which covered a period between 1979 to 2023.

However, since publications were not constant over time (**Fig. 1**), the sample size of years was *n* = 26. Each paper had on average 7.88 ± 4.91 statistical procedures (median = 7, range = 1 to 28 procedures, total = 261) and there was a significant increase in procedures (per paper) over the years **(Table 1, Fig. 1**).

**Table 1.** Generalized linear model showing which variables were related to statistical procedures described in papers. Values in bold indicate significant differences.

| Variables | $\chi^2$ | *P*-values |
| --- | --- | --- |
| Softwares | 38.01 | **0.0001** |
| Year | 9.67 | **0.0018** |
| Interaction | 2.32 | 0.5072 |

Computer softwares were first cited in papers dealing with plant-ant interactions in 2005, and since then 76% of papers (*n* = 108) reported the use of at least one software (mean per paper = 1.39 ± 0.76; median = 1; range 1 to 5; total number of softwares = 22; **Fig. 2a**). The R was initially adopted in plant-ant studies in 2012, and it has consistently been used even since, either alone or in combination with other softwares (**Fig. 2b)**. The use of R has increased over the years (*R^2^* = 0.69: *p* < 0.001); and it has been by far the most cited software (**Fig. 2a**). Papers that used R together with other softwares had significantly more statistical procedures than papers that used R alone (**Table 1, Fig. 1 and 2c**).

A total of 59 R packages were cited in the literature, of which *vegan* and *lme4* were used more frequently (**Fig. 2d**) (Online Resource). Out of the most used packages in plant-ant studies, five of them (*car, lme4, MASS, multcomp* and *vegan*) were also frequently used in biodiversity conservation, ecology and forestry (data in Lai et al., 2019; 2023a; 2023b). The pairwise comparisons according to the Jaccard index of dissimilarity yielded values of 0.76, 0.81 and 0.72 for the comparison between current data and biodiversity conservation, ecology and forestry, respectively. Nine of the top-used packages in plant-ant interactions (*bipartite, circular, DHARMa, glmmADMB, glmmTMB, maptools, stats, survival, survminer*) (**Fig. 2d**) were not found (as the top packages) in the other above-mentioned fields.

The GLMM is the most frequent test in plant-ant interaction studies, surpassing traditional tests such as *t*-tests and Anova (**Fig. 3**). The so-called modern tests have risen in frequency since their implementation, which coincides with the adoption of R software in 2012 (**Fig. 4**).

**Fig 4.**
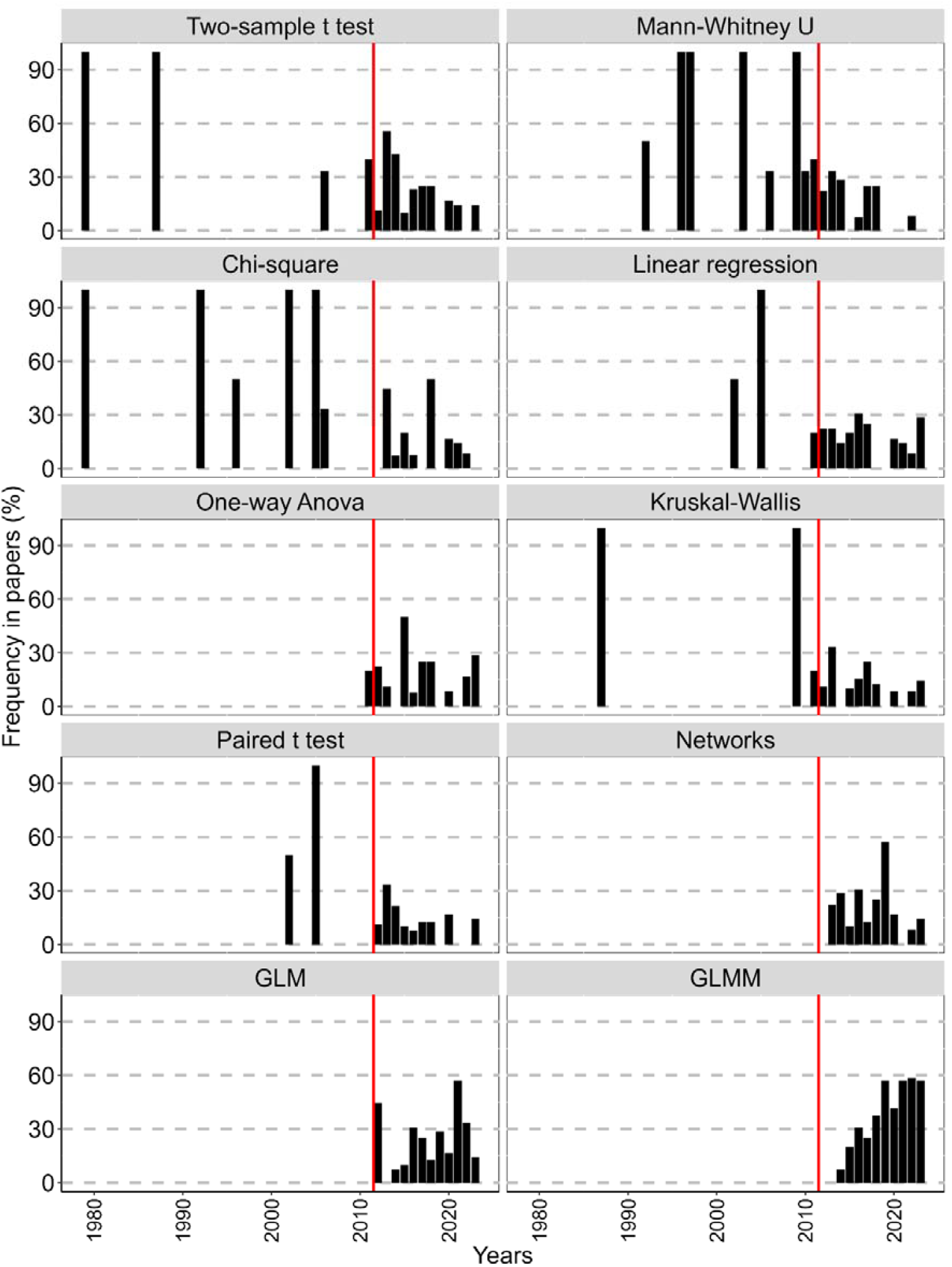
Temporal variation in the use of some analyses in plant-ant research. The vertical line marks the year of 2012, when R software was implemented in studies of plant-ant interactions.

## Discussion

This study demonstrates that scientists employ hundreds of different statistical procedures in research, and that the use of R software is associated with an increase in the number of statistical procedures over time. The use of traditional tests is declining as GLM(M)s and Network analyses become more consistently used in studies of plant-ant interactions. The R software is currently ubiquitous in literature, and the most used packages are associated with mixed models, community ecology and Networks. The packages used plant-ant interactions are quite different from those used in other fields.

Most statistical procedures were used only once or twice (∼ 60% of the dataset). This indicates that these procedures are applicable only to specific scenarios, and that they function primarily as prerequisites (e.g., outlier detection, normality tests) for leading tests. In contrast, the most frequently used procedures tended to be leading tests that generate *p*-values such as Anova, GL(M)M and *t* tests (Online Resource). GLMs, GLMMs and Networks were frequent in the dataset, surpassing traditional tests. This pattern indicates the growing importance of these approaches in ecology in recent years. Although such analyses appeared in 48% of all publications, they have been implemented only in 2012. This may imply that, for a considerable period, scientists relied on (and achieved successful outcomes with) a relatively small set of statistical techniques. However, modern analytical approaches enabled new study designs and the exploration of different types of data that cannot be adequately addressed using some classical tests.

With the widespread implementation of R software, handling large volumes of data is no longer a major challenge in ecology. R has been extensively adopted in ecological research and faces virtually no competition. Some procedures are designed to run exclusively in R; consequently, statistical literacy and basic programming skills have become essential (Zuur et al., 2010). Although ecologists began using R as early as 2008 (Gao et al., 2025), it appeared in Brazilian EFN–plant studies only in 2012. This four-year gap may reflect an initial period of adaptation to the software. A similar temporal pattern is observed in Mexican EFN–plant studies as well (see Díaz-Castelazo et al., 2013, 2010 for comparisons).

The use of R in plant–ant research is proportionally higher than in ecology overall (Gao et al., 2025).

Since its introduction into plant–ant studies in 2012, R has appeared in 79% of publications (compared with 42% in ecology broadly), and in 2023 it was cited in 85% of papers (versus 67% in ecology) (Gao et al., 2025). In three years (2019, 2021, 2023), R was used in all plant–ant studies examined. The strong prevalence of R indicates that its programming language has not posed a substantial barrier to researchers. To the best of my knowledge and experience, R remains the primary software for conducting GLMs and GLMMs. Network analyses can also be performed entirely within R (Barbosa et al., 2025; Câmara et al., 2018; Souza et al., 2024). Nonetheless, many authors complement R with additional programs to create figures or compute specific indices (e.g., Pajek, Modular, Aninhado) (Lange and Del-Claro, 2014; Miranda et al., 2019). The data indicate that the number of statistical procedures is significantly higher when researchers use R in combination with other software, which may perform simpler than R in some tasks.

In contrast with Gao et al. (2025), who reported *R^2^* values greater than 0.98 for the relationship between the use of R and years, the present study found a considerably lower *R^2^* (0.69). This difference arises from the temporal fluctuations in the adoption of R. For instance, although R appeared in more than half of the publications in its debut year (2012), its use declined during the subsequent three years, thereby reducing the strength of the regression.

The similarity between the most frequently used packages in plant–ant interaction studies and those from other disciplines (ecology, biodiversity conservation, and forestry) was generally low (Lai et al., 2023b, 2023a, 2019). The packages *vegan* and *lme4*, designed for community ecology and mixed models, respectively (Bates et al., 2025; Oksanen et al., 2025), were consistently prominent across all fields, indicating that certain analytical tools are pervasive and widely applicable. In the present dataset, half of the most frequently used packages were associated with linear (mixed) models (*lme4, nlme, emmeans, glmmTMB, DHARMa, glmmADMB, MASS, multcomp*), which show the ubiquity and necessity of these analyses in plant-ant studies.

A noteworthy package is *bipartite*, which is employed for Network analyses (Dormann et al., 2008). This approach has become increasingly common in plant–ant research (Lange et al., 2013; Miranda et al., 2019), enabling the investigation of multiple levels of community organization, core species, and interaction diversity (Dáttilo and Rico-Gray, 2018). The *bipartite* package, however, is rarely listed among the most widely used packages in ecology as a whole (Lai et al., 2019). This shows that, despite sharing some analyses with other fields (evidenced by the use of *vegan* and *lme4*), plant–ant interactions rely on a distinct subset of methodologies that require specialized analytical tools.

## Conclusion

Research on plant–ant interactions has undergone substantial changes in study design over the years, and statistical methods have rapidly evolved to meet the growing demand for more robust analyses and the integration of multiple variables. While statistics may play a relatively minor role in some areas of biology (Pekár and Brabec, 2016), they constitute a cornerstone of research in plant–animal interactions. Indeed, flawed analyses and misinterpretation of data are among the primary causes of manuscript rejection (Andrew, 2020). Consequently, ecologists cannot afford to apply inappropriate tests due to insufficient statistical knowledge (Fraker and Peacor, 2008). Within this context, the observed increase in statistical procedures over the years may reflect a natural progression of the field driven by statistical literacy and the demand for more robust analyses in studies of plant-ant interactions.

## Acknowledgments

Vanessa Dayane da Costa Barbosa, Rodrigo Augusto Santinelo Pereira, Alexandra Bächtold, and the staff and participants of the *Simpósio de Ecologia Comportamental e Interações*, 2025 - where this work was presented - for comments and suggestions.

## Author contribution information

The concept, writing, data collection and analysis of this research was made by the author

## Statements and Declarations

The author declares no competing or financial interests

## Data Availability Statement

Data is available as Supplementary materials (Online Resource) under request.

## Disclaimer

Artificial Intelligence tools were used to perform English proofreading only.

